# Environmental and Population influences on Mummichog (*Fundulus heteroclitus*) Gut Microbiomes

**DOI:** 10.1101/2024.01.26.577513

**Authors:** Lei Ma, Mark E. Hahn, Sibel I. Karchner, Diane Nacci, Bryan W. Clark, Amy Apprill

## Abstract

The mummichog, *Fundulus heteroclitus*, an abundant estuarine fish broadly distributed along the eastern coast of North America, has repeatedly evolved tolerance to otherwise lethal levels of aromatic hydrocarbon exposure. This tolerance is linked to reduced activation of the aryl hydrocarbon receptor (AHR) signaling pathway. In other animals, the AHR has been shown to influence the gastrointestinal-associated microbial community, or gut microbiome, particularly when activated by the model toxic pollutant 3,3’,4,4’,5-pentachlorobiphenyl (PCB-126) and other dioxin-like compounds. In order to understand host population and PCB-126 exposure effects on mummichog gut microbiota, we sampled two populations of wild fish, one from a PCB-contaminated environment (New Bedford Harbor, MA) and the other from a non-polluted location (Scorton Creek, MA), as well as laboratory-reared F2 generation fish originating from each of these populations. We examined the bacteria and archaea associated with the gut of these fish using amplicon sequencing of small subunit ribosomal RNA genes. Fish living in the PCB-polluted site had high microbial alpha and beta diversity and an altered microbial network structure compared to fish from the non-polluted site. These differences between wild fish were not present in laboratory-reared F2 fish that originated from the same populations. Microbial compositional differences existed between the wild and lab-reared fish, with the wild fish dominated by Vibrionaceae and the lab-reared fish by Enterococceae. These results suggest that mummichog habitat and/or environmental conditions has a stronger influence on the mummichog gut microbiome compared to population or hereditary-based influences. Mummichog are important eco-evolutionary model organisms; this work reveals their importance for exploring host-environmental-microbiome dynamics.

## Introduction

Microbial communities are central to animal biology, and the animal host plus its symbiotic microbial community is increasingly considered a single complex assemblage, termed the “holobiont”[1,2]. The microbiome can influence critical host processes including physiology, development, and behavior, which in turn contributes to higher order phenomena including host adaptation, co-evolution of host and microbiome, and genetic divergence[3]. As animals adapt to human-altered environments, it is important to incorporate host-microbe interactions as a factor affecting host ecology and evolution.

Fish are valuable field and laboratory models for investigating vertebrate host-microbiome interactions[4]. In fish, gut microbiomes influence a variety of host processes, including the digestion of plant material[5], immune regulation[6], and neurological development[7]. Unlike terrestrial mammals, fish live in a dense aquatic microbial environment that may interact with and influence the host microbiome. Changing environmental conditions such as pollution can cause rapid and profound shifts in environmental microorganisms, as well as aquatic host genetics and microbiome structure and function. Recent research has shown that external variables such as season, diet, and population density as well as host sex and immune system variation contribute to changes in fish gut microbiome richness and taxonomic composition (reviewed in [8]). A key research question is whether the composition of the microbiome is affected primarily by external environmental factors or by the host’s physiology. Answering such questions in fish requires attention to the host, its microbiome, and that of the environment[9].

The mummichog (*Fundulus heteroclitus*; also known as Atlantic killifish) is a unique natural model system for interdisciplinary study of the effect of host genetics (and the adaptation of the host to a particular environment) on gut microbiome assembly and function in relation to different environments. For decades, this estuarine fish has served as a mature model system for studies in ecophysiology, evolution, and immunology[10]. These fish possess a high standing genetic diversity, which allows them to adapt to changing environmental conditions, including pollution, salinity, and eutrophication. Because of their broad distribution across a wide range of environmental conditions, mummichog have been proposed as a teleost model for environmental effects monitoring[10,11]. A better understanding of mummichog microbiomes will improve their utility as an ecosystem indicator and as a model organism for investigating host-microbe interactions in changing environments.

Currently, little is known about the natural variation in mummichog microbiota and how it has changed following host adaptation to environmental shifts, in particular to increased pollution. Multiple waterways on the Atlantic coast are impacted by high levels of pollution in the form of polycyclic aromatic hydrocarbons (PAHs), chlorinated dibenzo-p-dioxins, and polychlorinated biphenyls (PCBs). Mummichog in these locations have experienced rapid and parallel adaptation to tolerate otherwise lethal concentrations of these chemicals[12]. The mechanism of aromatic hydrocarbon tolerance in mummichog involves selection for downregulation of the aryl hydrocarbon receptor (AHR) pathway [12–14]. In addition to affecting host genetics, environmental chemical contaminants including aromatic hydrocarbons may also shift the mummichog gut microbiome directly by altering microbial growth or indirectly by affecting host physiology [for a review see [15]].

Adaptation to polluted environments through altered function of AHR pathway genes may contribute to changes in the gut microbiota of mummichog. The AHR is prominent in regulating intestinal immunity[16] and has been shown in a variety of animals to influence the gut microbiome. In mice, experiments have shown that AHR contributes to the community structure of the cecal microbiota[17–19] while exposure to the potent AHR agonist PCB-126 increased inflammation and disrupts the gut microbiota, although the role of AHR was not directly investigated[20]. In zebrafish, AHR activation by PCB-126 is associated with altered growth of gut microbes and inflammation[21]. Examining the microbiome of PCB-exposed and PCB-adapted mummichog can help us further understand the consequences of pollution for the holobiont.

Here we compared the microbiomes of mummichog originating from Scorton Creek and New Bedford Harbor. New Bedford Harbor mummichog have adapted to tolerate high levels of PCB pollution in the sediment as a result of decades of industrial waste dumping, while the Scorton Creek mummichog population is sensitive to PCB exposure[12,22]. We examined gut microbiomes of these two mummichog populations taken directly from the wild, as well as after being reared for two generations (3-5 years) in a common garden lab environment. We hypothesized that the tolerant wild mummichog microbiomes would be distinct from the sensitive wild mummichog microbiomes and that this effect would persist to some degree in the lab environment. In particular, we expected the tolerant mummichog gut microbiomes to show signatures of dysbiosis, such as pathobiont expansion or increased beta diversity.

## Materials and Methods

### Field sampling

Wild mummichog were collected from Scorton Creek, MA (GPS: 41.746178, -70.426606) and New Bedford Harbor, MA (GPS: 41.657413, -70.918928) on September 9th, 2020 and September 22, 2020, respectively. Forty adult males were collected from each site using minnow traps placed at shallow depths (0.5-1.0 m) and transported live in a cooler with aerated water to Woods Hole, MA. Replicate 2-L surface water samples were collected from each site and filtered through 0.22 µm pore size Supor filters (25 mm; Pall Corporation) and frozen in cryovials at -80°C. Fish were sedated in 1.5% MS-222 and sodium bicarbonate and sacrificed by severing the spinal cord. The progenitors of the F2 fish were collected from Scorton Creek and New Bedford Harbor in the years 2016-2018 and were maintained for two generations separately in otherwise similar tanks with continuous flow through seawater at the Atlantic Coastal Environmental Science Division as described previously [23]. F2 fish experienced the same seawater and feeding conditions and can be considered common gardened. A mixture of male and female F2 fish from each originating population (n = 15 Scorton Creek F2; n = 44 New Bedford Harbor F2) were collected in June 16, 2021, flash frozen and shipped to Woods Hole, MA, where they were thawed and dissected. Aquaria water samples were not available for comparison.

### Sample processing and sequencing

Prior to dissection, measurements of total length and body weight of each fish were taken. Then, the entire digestive tract from the esophagus to the anus was removed using sterile tweezers and frozen at -80°C. DNA was extracted from the guts and filters using the DNEasy PowerBiofilm kit (Qiagen) according to manufacturer instructions. Eight DNA extraction controls, consisting of no sample, were processed along with the biological samples. Barcoded primers 515FY[24] and 806RB[25] were used to amplify the V4 region of the small subunit rRNA gene in bacteria and archaea. 1 µL of DNA template was included in a 25 ul GoTaq Flexi PCR reaction. Two PCR controls consisting of 1 µL of PCR-grade water as template were also included, as well as microbial genomic DNA from a Human Microbiome Project mock community (BEI Resources, NIAID, NIH as part of the Human Microbiome Project: Genomic DNA from Microbial Mock Community B (Even, Low Concentration), v5.1L, for 16S RNA Gene Sequencing, HM-782D). PCR conditions were as follows: 37 cycles (95°C 20s, 55°C 15s, 72°C 30s) with a 2 minute 95°C hot start and 10 minute 72°C final elongation. Each reaction was run in duplicate and then pooled for gel purification (MinElute PCR Purification Kit, Qiagen). Purified products were quantified using the Qubit 2.0 fluorometer HS dsDNA assay (ThermoFisher Scientific), diluted to equal concentrations, and pooled for sequencing. Sequencing was performed at the University of Georgia Genomics and Bioinformatics core on a paired-end 2×250 bp Illumina MiSeq platform.

### Data analysis

All code used for generating the data and figures for this project can be found on GitHub: https://github.com/microlei/killifish_manuscript. All raw sequence data have been uploaded to the Sequence Read Archive under BioProject ID PRJNA858104. Raw reads of the V4 region of the 16S rRNA gene were processed using the DADA2 package in R (v. 3.6.2)[26]. Reads were filtered using the default parameters of the function filterAndTrim except both forward and reverse reads were truncated at 240 bp. Read merging, chimera removal, and amplicon sequence variant (ASV) generation was also performed using DADA2. Taxonomy was assigned at the 100% identity level based on the Silva SSU rRNA gene database (v. 138)[27]. Contaminant reads were identified and removed, using the negative controls as a guide, by the prevalence method of the R package decontam[28]. Sequences belonging to the Kingdom Eukaryota or Order Chloroplast were manually removed. Data analysis was performed in RStudio using the packages vegan and phyloseq[29,30]. Alpha diversity metrics were estimated using the function estimate_richness in vegan on the raw read counts. Differences in alpha diversity were tested using the pairwise t-tests with holm correction for multiple comparisons[31]. Counts were transformed to the center log ratio after removing zeros using the package zCompositions[32]. Center log ratio (CLR) transformed counts were used for the Principal Component Analysis (PCA) and Aitchison distances (the Euclidean distance between the CLR transformed counts) used for distance measures between samples. Aitchison distances were used because it is robust to subsetting of the data and accounts for the compositionality of relative abundance data[33]. Differentially abundant taxa were identified using the corncob package[34]. Microbial interaction networks were generated in the R package SpiecEasi[35], using the graphical lasso method. Network metrics were calculated and analyzed using igraph[36]. For calculating average path length, only the lengths of the existing paths were considered and averaged. Networks were visualized using ggraph[37]. To reduce computational time, taxa from the wild gut microbiome samples were pruned for taxa with at least 50 reads total across all samples, but the sequences from the F2 microbiomes were not pruned due to their lower diversity. Another network was generated for New Bedford Harbor wild, pruning a greater proportion of lower abundance taxa to generate a network with an identical number of taxa as the Scorton Creek wild network to test whether the results were a consequence of the higher number of nodes.

## Results

### Sequencing statistics

After quality control and filtering, a total of 12,129,065 reads were retained from the 139 fish gut microbiome and the 4 water microbiome samples. Some fish gut microbiome samples had a high proportion of chlorophyll reads but the median number of reads per sample was 88,125 with a range of 6,412 to 214,684 reads per sample. A total of 11,132 Amplicon Sequence Variants (ASVs) were identified, with a median of 83 unique ASVs per fish.

### Alpha and beta diversity

Gut microbiome alpha diversity, including richness, Shannon index, and Simpson index, differed between the two wild populations but not between the F2 fish populations (Figure 1A, pairwise t-test: p<0.0001 for all significant comparisons). The number of unique taxa (Richness) in New Bedford Harbor (NBH) wild fish (the tolerant population) was on average 412 with standard deviation (sd) of 200, which was similar to that of the water samples (552 taxa, average of 2 samples). In contrast, the alpha diversity observed in Scorton Creek (SC) wild fish (the sensitive population) (96, sd = 112) and all F2 fish populations (82, sd = 47) was lower, while that of the water from SC was similar to NBH water (478, average of 2 samples) (Table S1). Similarly, the NBH wild fish had a more even distribution of microbial taxa (Shannon index) than the other three fish populations. Overall, pairwise t-tests show that NBH wild fish have significantly different diversity values than all other fish types while SC wild, SC F2, and NBH F2 fish have similar measurements (Table S5). The Simpson index, a measure of the uniformity of the species abundances, showed similar patterns. One exception is that SC wild fish had a higher Simpson index (0.439) than NBH F2 fish (0.282). Richness of water samples collected at the wild sites was higher compared to SC but not NBH wild fish gut microbiota (Figure 1A).

**Figure 1.**
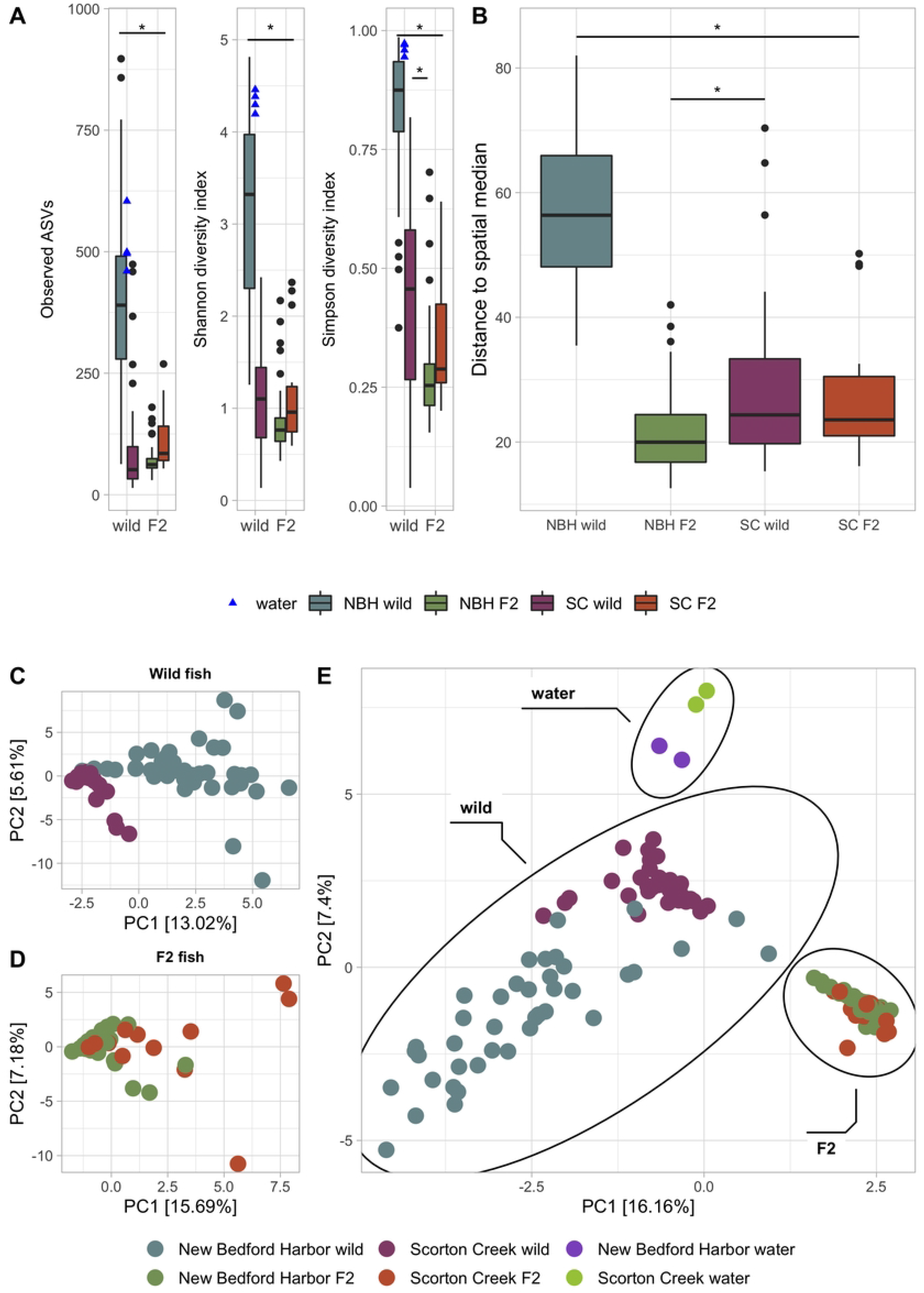
Alpha and beta diversity measurements of mummichog microbiomes. (A) Alpha diversity indices. Asterisk indicates significant differences (p<0.05) by pairwise t-test with Holm correction between NBH wild vs all other fish and between SC wild and NBH F2. (B) Beta diversity calculated as distance to spatial median in Aitchison distance. Asterisk indicates significant differences (p<0.05) by Tukey HSD pairwise testing. (C-E) PCA plots based on Aitchison distances of wild fish (C), F2 fish (D) and all samples (E). Ellipses around water samples, wild fish microbiomes, and F2 fish microbiomes are drawn to visually highlight the groups.

For examining beta diversity, a Principal Component Analysis (PCA) of the Aitchison distance comparing microbiomes showed that water, as well as gut microbiomes of wild fish and F2 fish were distinct (Figure 1E). The microbiomes of the two wild fish populations separate from each other while the two F2 fish populations were not as distinct. Separate PCAs of just wild or F2 fish show more separation between the NBH and SC populations (Figure 1C and 1D).

PERMANOVA comparisons of the Aitchison distances between the four fish types (SC wild, NBH wild, SC F2, and NBH F2) demonstrates that each fish type is significantly different from the others, as are the wild fish and F2 fish. However, the percent of variance explained (R2) varies between comparisons (Table 1).

**Table 1.** PERMANOVA results comparing microbiome composition between fish types.

| Comparison | Df | Sum Sq | R<sup>2</sup> | Pseudo-F | p-value |
| --- | --- | --- | --- | --- | --- |
| **All fish types** | 3 | 92207 | 0.302 | 19.5 | 0.001 |
| **Wild vs F2** | 1 | 62184 | 0.204 | 35.0 | 0.001 |
| **SC wild vs NBH wild** | 1 | 27072 | 0.134 | 12.1 | 0.001 |
| **SC F2 vs NBH F2** | 1 | 2878 | 0.073 | 4.5 | 0.001 |

Each row is an independent PERMANOVA test comparing the Aitchison distances of the mummichog microbiomes. All results were significant, although the R<sup>2</sup> value was variable. NBH, New Bedford Harbor; SC, Scorton Creek; Df, degrees of freedom; R<sup>2</sup>, R-squared value; Pseudo-F values derived from 999 permutations.

Comparison of beta dispersion showed that the NBH wild population had a significantly higher beta diversity compared to all other fish types (Tukey HSD p< 0.001) and SC wild fish had a higher beta diversity than NBH F2 (Tukey HSD p< 0.05) (Figure 1B). The F2 fish microbiomes did not differ in beta diversity (Tukey HSD p > 0.05). After accounting for the effect of fish type, fish length, weight, and (in the case of F2 fish) sex were not significant factors in explaining the variance of Aitchison distances across samples (Table S2).

### Core and differentially abundant taxa

To identify the core gut microbiome of the mummichog, we looked at the shared taxa between populations. The two wild populations shared 563 ASVs (6.3% of the total wild microbiome), while the two F2 populations shared 283 ASVs (19.5% of the total F2 microbiome). A total of 53 taxa were shared between all fish types. In general, there was a low percentage of overlap in ASVs between fish types, even between common gardened F2 fish (Figure 2A). The taxa shared between all fish types included members of the Phyla Firmicutes, Proteobacteria, and Planctomycetota. Detailed taxonomic classification of these shared ASVs can be found in Table S3.

**Figure 2.**
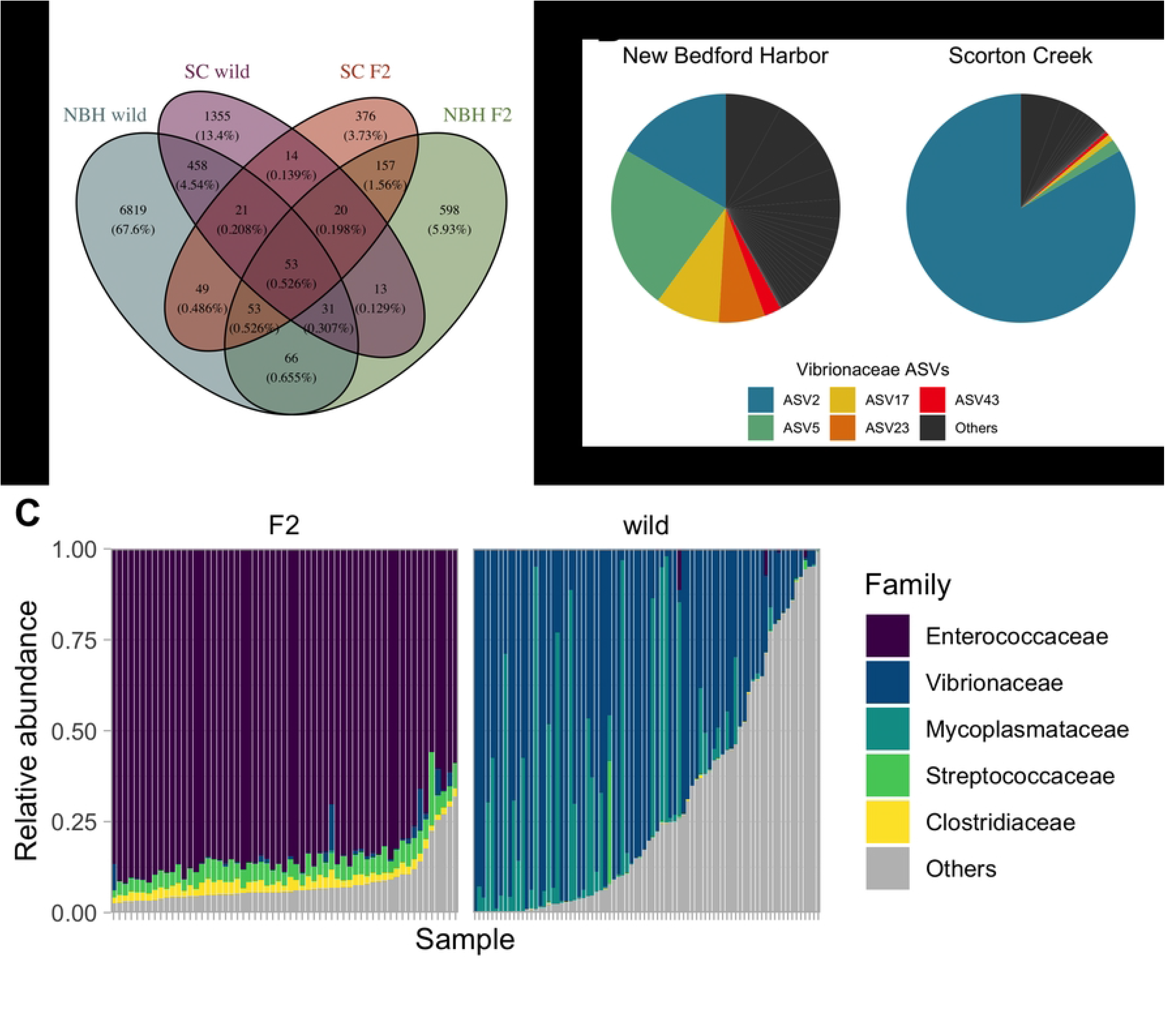
Core and differentially abundant Amplicon Sequence Variants (ASVs) across fish populations. (A) A Venn diagram showing the overlap in ASVs between fish types, with 53 ASVs (0.526% of ASVs) shared among all four fish types. SC = Scorton Creek. NBH = New Bedford Harbor. (B) Pie chart of ASVs from the family Vibrionaceae shared between the two wild fish populations. Chart area is proportional to relative abundance. Colored ASVs have significantly different abundances between NBH wild and SC wild. (C) Bar graph of the relative abundances of selected bacterial families from all mummichog gut microbiomes. Colored taxa are the families of ASVs identified as significantly different between F2 and wild fish microbiomes.

Differential abundance (DA) analysis was performed on subsets of the dataset to investigate which taxa may be driving the differences in community composition. In a comparison between all wild fish and all F2 fish gut microbiomes, DA analysis found 37 significant taxa. Overall, the dominant taxa in the wild fish were from the families Vibrionaceae or Mycoplasmataceae, while the dominant families of the F2 fish were comprised of Enterococcacea, Streptococcacae, and Clostridiacea (Figure 2C).

In the wild fish gut microbiomes, 40 taxa were significantly differentially abundant between SC and NBH fish - that is, enriched in either SC or NBH guts. We examined the relative abundance of these taxa in the seawater samples and found that they were at very low abundance (<1%) or not present at all. Of those that were present, only 12 out of 28 taxa had a higher abundance in the water sample corresponding to the enrichment in the gut samples (Table S4). Within wild fish, SC guts were characterized by high levels of ASV2, an unclassified Vibrionaceae, while NBH guts had a more even distribution of several Vibrionaceae ASV relative abundances (Figure 2B). Only 3 taxa were identified as significantly differentially abundant between the two F2 populations: the unclassified Vibrionaceae (ASV2), a member of the *Paracoccus* genus, and member of the *Colwellia* genus. Of note, ASV2 was significantly enriched in both wild and F2 SC fish compared to their NBH counterparts.

### Network analysis

Visualization of the networks generated for each fish type revealed that the NBH wild fish microbiome network consisted of one large connected graph while the other three networks were comprised of multiple disconnected subgraphs as well as numerous isolated taxa (Figure 3A). The degree distribution for NBH wild was also distinct from the others, showing a humped shape rather than an exponentially declining shape (Figure 3B). Network characteristics were calculated for each graph and are summarized in Table 2. Because of the disconnected nature of three of the graphs, many common centrality measures are not defined and thus not included. Compared to SC wild and NBH F2, NBH wild fish displayed lower transitivity (also known as clustering coefficient), longer average path length, lowered modularity, and lowered degree centralization. The F2 fish displayed higher edge density and degree centrality than the wild fish and lower average path lengths. Regenerating the NBH wild network by subsetting to a number of abundant taxa equal to the number of nodes in the SC wild network did not substantially change its network characteristics (Figure S1 and Table 2). Network stability analysis revealed that the connectivity of the SC wild network was more sensitive to node removal than the NBH wild network (Figure S3).

**Figure 3.**
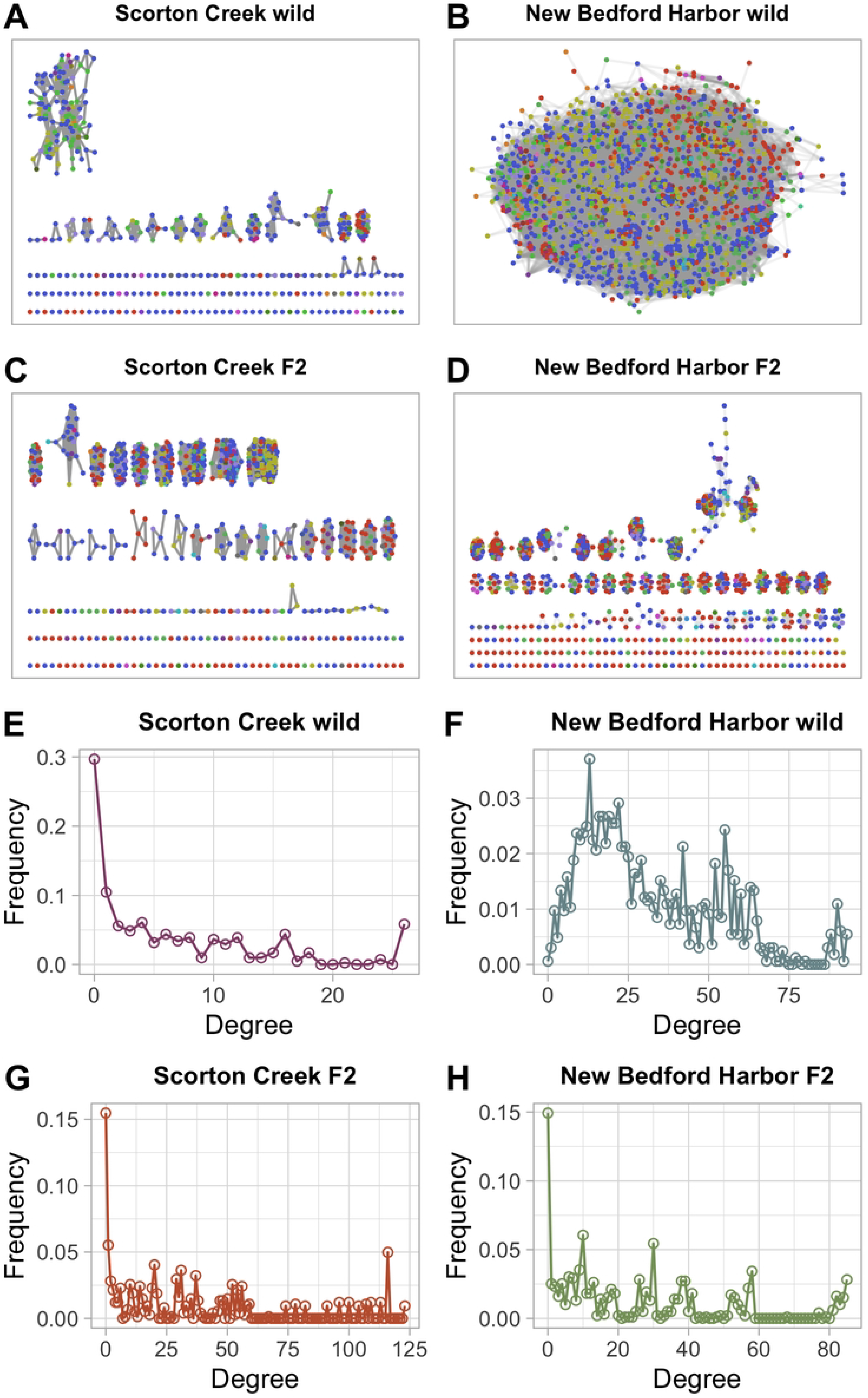
Visualization of the microbial interaction networks of each fish type. (A-D) Comparison of microbial networks show that the New Bedford Harbor wild network is fully connected while the other three networks are comprised of multiple disconnected components and many isolated ASVs. Nodes represent ASVs, and edges between nodes represent associations. Nodes are colored by Phylum affiliation. (E-H) Degree distributions of the networks shown in A-D demonstrate that the NBH wild degree distribution is hump shaped, while the other networks have a decaying degree distribution. This indicates that NBH wild network is more similar in structure to a random network.

**Table 2.**
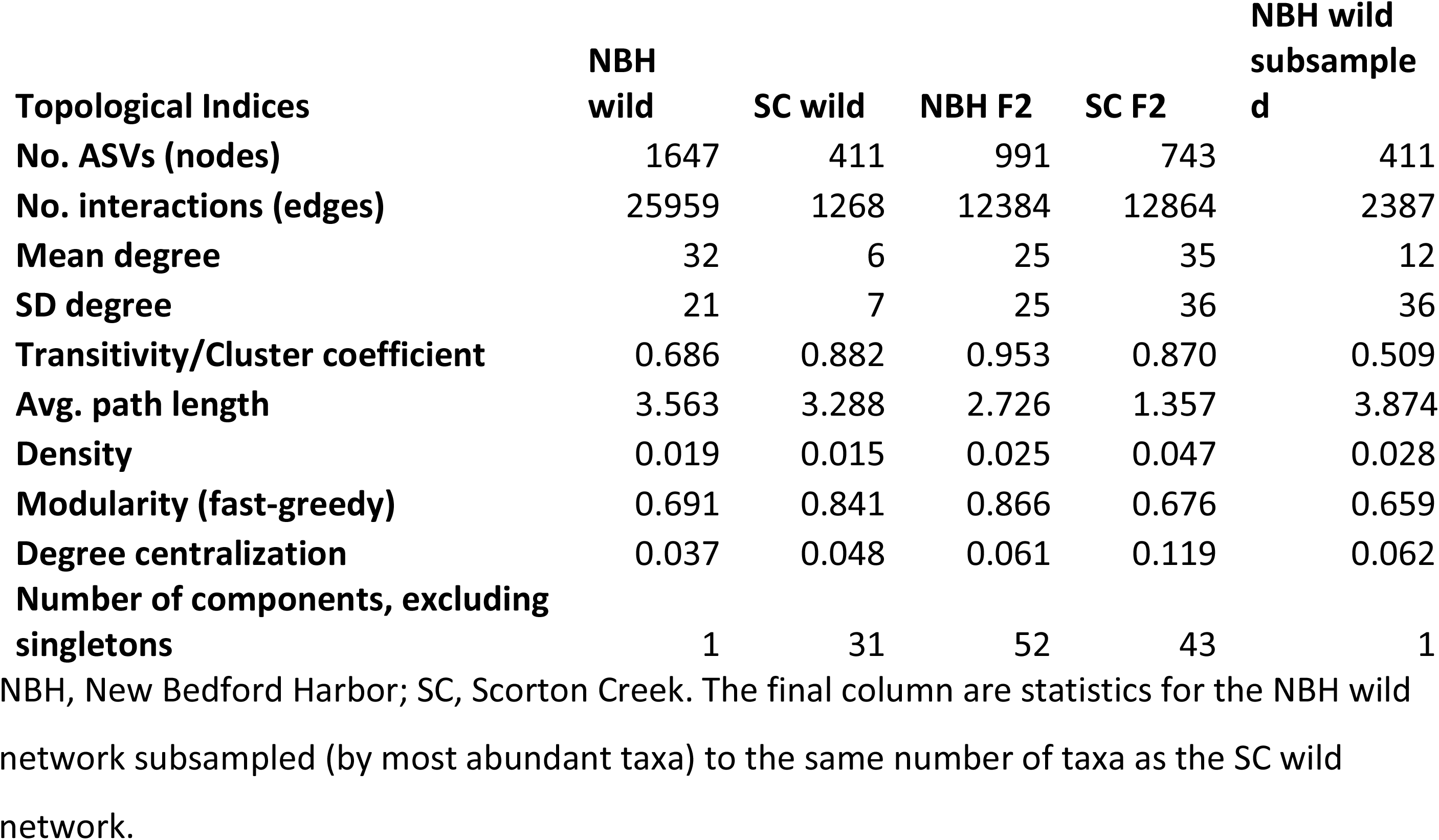
Network topology metrics for the microbiome networks.

Some SC F2 network properties were markedly different from that of the NBH F2 network, despite both fish populations living in a common garden environment. The SC F2 network had a higher edge density and lower shortest path length than NBH F2, despite being overall less densely connected (transitivity). However, SC F2 network was generated using 15 samples while the NBH F2 network was generated using 44 samples, which may affect the quality of associations inferred.

## Discussion

In this study, we examined and compared the gut microbiomes of wild and lab reared mummichog from two populations distinguished by their differential sensitivity to aromatic hydrocarbon pollution. We found that alpha diversity, beta diversity and overall community composition of mummichog gut microbiomes are highly plastic and are strongly influenced by the environment, although host population (fish type) also has some effect. Mummichog size, weight, and sex were not correlated with the microbiome composition. The microbiomes from wild, PCB-resistant NBH fish showed potential signs of dysbiosis characterized by high beta diversity and altered network structure compared to the other fish types. Despite the large differences in dominant taxa between captive and wild fish, network analysis revealed very similar network structure in the microbiomes of all fish types living in non-PCB contaminated environments.

### Mummichog microbiomes are dynamic and highly influenced by their environment

The composition of the gut microbiomes of all four fish types were distinguishable from each other (Figure 1C-E), suggesting that habitat/environment as well as population influence the gut microbiome. However, high microbial diversity in the surrounding aquatic habitat does not necessarily lead to high gut microbiome diversity, as evidenced by the difference in alpha diversity metrics between the SC wild fish and its water source. This suggests that the mummichog gut microbiome is a selective environment, likely due to a combination of host control and environmental filtering. This result is in line with previous work describing fish gut microbiomes as specialized environments with potentially co-evolved relationships between host and gut bacteria[38–40]. Variation in microbial composition between populations of the same species of fish have been explained by a combination of environmental and host effects. For example, in three-spined stickleback, host genetic distance correlated with the microbiome’s UniFrac distance while the microbiota also resembled that of local invertebrate prey[41]. In a study comparing inter- and intra-species microbiome differences of Norwegian cod fishes, niche separation, and to a lesser extent evolutionary separation, were major contributors to microbiome differences[42]. Mummichog gut microbiomes have also been found to differ both between wild populations and between wild and captive raised fish[43,44]. The wild mummichog in the present study experienced two different environments (New Bedford Harbor, Scorton Creek) and previous work on these populations has shown them to be genetically distinct[45], suggesting that environmental and host immune differences may contribute to differences in gut microbiomes. All fish were sampled in the summer to reduce the effect of seasonality, but dietary input, water condition, and other features likely vary between the two wild sites. Additionally, NBH fish were exposed to high levels of PCB and likely other industrial pollutants due to its more urban location. Exposure to environmental chemical contaminants has been shown in multiple fish species to be disruptive to gut microbiome composition[46–48]. Upon comparing the variation in microbiome composition between the two wild fish populations and the common gardened F2 fish, we suggest that the environment is likely the dominant force driving differences in mummichog gut microbiome composition in these populations.

In addition to environmental influences, our comparison between F2 fish populations reveals the potential for host effects on the gut microbiome. The microbial community composition of SC F2 and NBH F2 fish were significantly different as assessed by PERMANOVA. However, the effect size was small, and the result may also reflect the differences in sample size (NBH F2 consisted of 44 fish while SC F2 consisted of 15 fish). Differential abundance analysis revealed another indicator of host effect: ASV2, an unclassified member of the Vibrionaceae family, is significantly enriched in both the wild and F2 SC population compared to their NBH counterparts. Although Vibrionaceae is not dominant in F2 fish generally, its abundance in both SC fish types may be a result of host selection. Within fish populations, genetic factors such as MHC genotype in stickleback and Bacterial Cold Water Disease resistance in rainbow trout have been shown to affect microbiome composition[49,50]. Closer examination of the genetic and microbial differences of these common gardened captive mummichog populations could reveal more concrete links between host genotype and microbiome composition.

### Residing in polluted water may impact both microbiome composition and structure

Our most striking finding was the increased diversity and difference in network structure of the NBH wild fish microbiomes compared to the three other fish types. NBH wild was the only fish population that was directly exposed to PCBs. Although the fish are apparently healthy due to their tolerant phenotype, their microbiomes suggest a difference in host-microbiome interactions.

The increased beta diversity of NBH wild microbiomes is in line with the *Anna Karenina Principle* of dysbiosis, in which external environmental stress alters the microbiome in unpredictable ways, increasing variation among individuals[51]. The resulting high beta diversity may be due to direct perturbation of the microbiome, such as by displacing mutualists, or due to a compromised host immunity. Laboratory-raised F2 generation NBH fish were previously found to have no difference in susceptibility to *Vibrio harveyi* infection compared to lab-reared PCB-sensitive fish from a reference site, but no comparison was made between lab-reared and wild-caught fish[23]. Without additional experimental evidence, it is not possible to determine whether NBH wild fish display increased tolerance to pathogens than their captive counterparts or whether the increase in gut diversity is related to disease at all. The increased alpha and beta diversity in NBH wild fish is consistent with the hypothesis that NBH wild fish may have a reduced ability to regulate its gut microbiome. On the other hand, low gut alpha diversity and stability in fish is commonly associated with disease[52]. In the case of mummichog, it is evident that diversity alone is not sufficient to infer dysbiosis as both increases and decreases in diversity can be implicated in gut microbiome disorders[53].

A healthy gut microbial community provides necessary services to the host, including the degradation of fiber and production of metabolic products, contributing to host nutrient absorption and energy metabolism[54]. These metabolic functions are distributed among microbial consortia in the gut and therefore the redundancy and stability of microbe-microbe interactions is a topic of interest for those studying gut microbiomes[55].

Both theoretical and experimental studies have highlighted the need to understand the interaction network of microbial communities as it relates to microbiome functioning[46,56,57]. Given the presumption of altered host regulation in the NBH wild fish microbiomes, we also expected to find a difference in the network structure of the microbial interactome. Specifically, we expected the network to be less densely connected and to display with less structure.

Network analysis revealed that the NBH wild microbiome is indeed structured differently than the other three fish types. The network is larger, consisting of over 1000 nodes, and the edges appear to be arranged more randomly (Figure 3). In contrast to our expectations, the NBH wild network is one large connected unit while the other networks consist of multiple disconnected components and isolated nodes. The disconnected nature of the SC wild and the F2 networks is surprising because loss of connectivity is associated with impaired gut microbiome health and such a microbiome is predicted to perform poorly in its services to the holobiont[56].

A closer examination of the network properties suggests that the NBH wild network had lower overall density and a reduced structure, in line with our expectations. Transitivity, which quantifies the degree to which nodes cluster together, and modularity, which quantifies the extent to which the network is comprised of tightly connected components, were both low in the NBH wild network (Table 2). Additionally, the network degree distribution of NBH wild fish is more similar to that of a random network topology than that of a scale-free or small world network typically found in microbial communities (Figure 3B). Microbial networks typically have small average shortest path lengths[58], but the NBH wild network has the longest path length of the three network types. The observed differences in network properties were not due to the comparatively large size of the NBH wild network. When the NBH wild dataset was subset to the same number of taxa as the SC wild fish, the same overall patterns emerged (Figure S2 and Table 2).

Overall, the NBH wild microbiome network, while fully connected, appears to be less structured internally than the disconnected networks of the other three fish types. Combined with the differences in diversity, these results support the hypothesis that the NBH wild microbiome is perturbed in some way as a result of its host population’s aquatic environment. In addition to being highly polluted, the New Bedford Harbor site is low energy, more urban, and has greater turbidity and higher organic content compared to the swifter waters of Scorton Creek

### Captive fish microbiomes are compositionally distinct from, but have interaction networks similar to, their wild counterparts

While the wild fish gut microbiomes were dominated by Proteobacteria, the F2 fish microbiomes were dominated by Firmicutes, including the highly abundant *Enterococcus* ASV1. This dramatic shift in microbiome composition from wild to captive reared fish has been documented in aquaculture. Captive rearing of fish alters many factors which have been shown to influence gut microbiota, including stocking density, salinity, diet, and antibiotic treatment[39,40]. For example, the gut microbiota of rainbow darter (*Etheostoma caeruleum*) decreased in alpha diversity upon acclimation to a laboratory environment, and the dominant taxa shifted from Proteobacteria and Firmicutes to mainly Firmicutes[59]. The beta diversity of the acclimated fish microbiomes was also affected. In contrast, the microbiomes of hatchery-born Atlantic salmon juveniles displayed a higher alpha diversity than wild-born salmon while wild salmon hosted a more specialized microbial community[60]. Additionally, the commercial diet fed to these salmon contributed to the dominance of the Lactobacillaceae family, which was absent from the abundant taxa of the wild microbiomes.

A complete turnover of the microbial environment can have consequences for the physiology of the host. Exposure to different microbes through the life of a fish activates a multitude of immune signaling pathways and commensal microbes can modulate systemic immune responses in the host[61]. Thus, we may expect host regulation of gut microbiota in the lab-reared mummichog to differ from that of wild mummichog. Such a difference would restrict the application of inferred host-microbiome interactions in a controlled lab setting to wild fish. However, despite the large differences in composition, the network structure of the F2 fish gut microbiomes was very similar to that found in the guts sampled from SC wild fish. This result indicates that the disconnected, multi-component network may be a conserved feature of mummichog microbiomes, and that host regulation of intestinal microbiota may function similarly between wild and F2 fish. The densely connected components may represent functional modules that are maintained by the host. However, without further sequencing of the microbial functional genes, it is difficult to come to any conclusions about the overall metabolic potential represented by the taxa present in each component. Additionally, information on host transcriptional activity in the gut environment is needed to confirm any similarity in intestinal regulation.

## Caveats and conclusions

Because only one tolerant and one sensitive population was sampled, the generalizability of these results is limited. We do not know what specific environmental conditions have contributed to the differences in these microbiomes. Additionally, direct host-microbiome interactions could not be assessed without experimental manipulations. Future work on the microbiomes of these and other populations of mummichog should ideally incorporate an experimental component as well as examine the functional profile of both microbiome and host.

In conclusion, this study establishes a baseline understanding of the microbiome composition of wild and captive mummichog and provides evidence for a link between acquiring a divergent microbiome and living in a PCB-contaminated environment. We were able to describe the differences in microbiome composition and structure in the PCB-exposed mummichog population compared to the other fish populations that were not exposed to PCBs. We discovered that the dominant phyla of the mummichog microbiome can switch after two generations of captivity, although the structure of the microbe-microbe interactions remains similar. Finally, the plasticity of the mummichog microbiome highlights the strong connection that fish microbiomes have to their environment as well as the need for further research on the consequences of environmental change on the holobiont.

## Acknowledgements

We thank Sara Hu, Jacob Cram, and Samantha Gleich for valuable statistical input, Neel Aluru and Carolyn Miller for technical support, and Ian Kirby, Joe Bishop, Madison Francoeur, Hannah Schrader, and Tara Burke for fish care and breeding.

## Supporting information

**Table S1.**
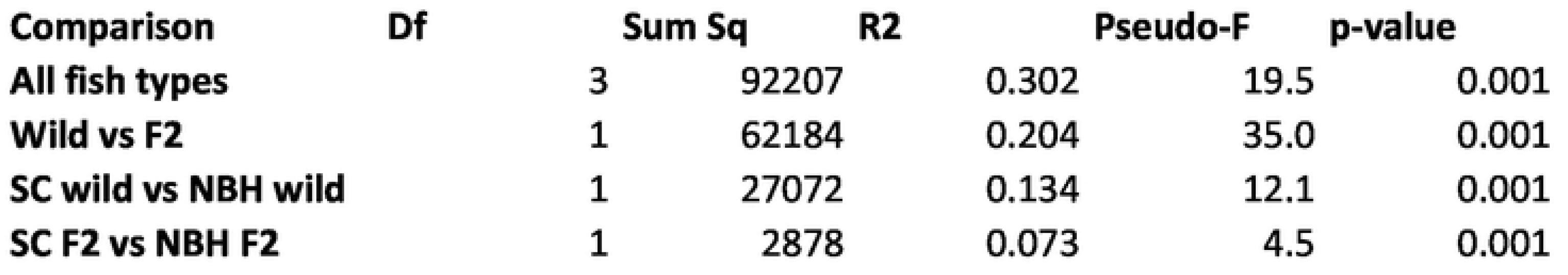
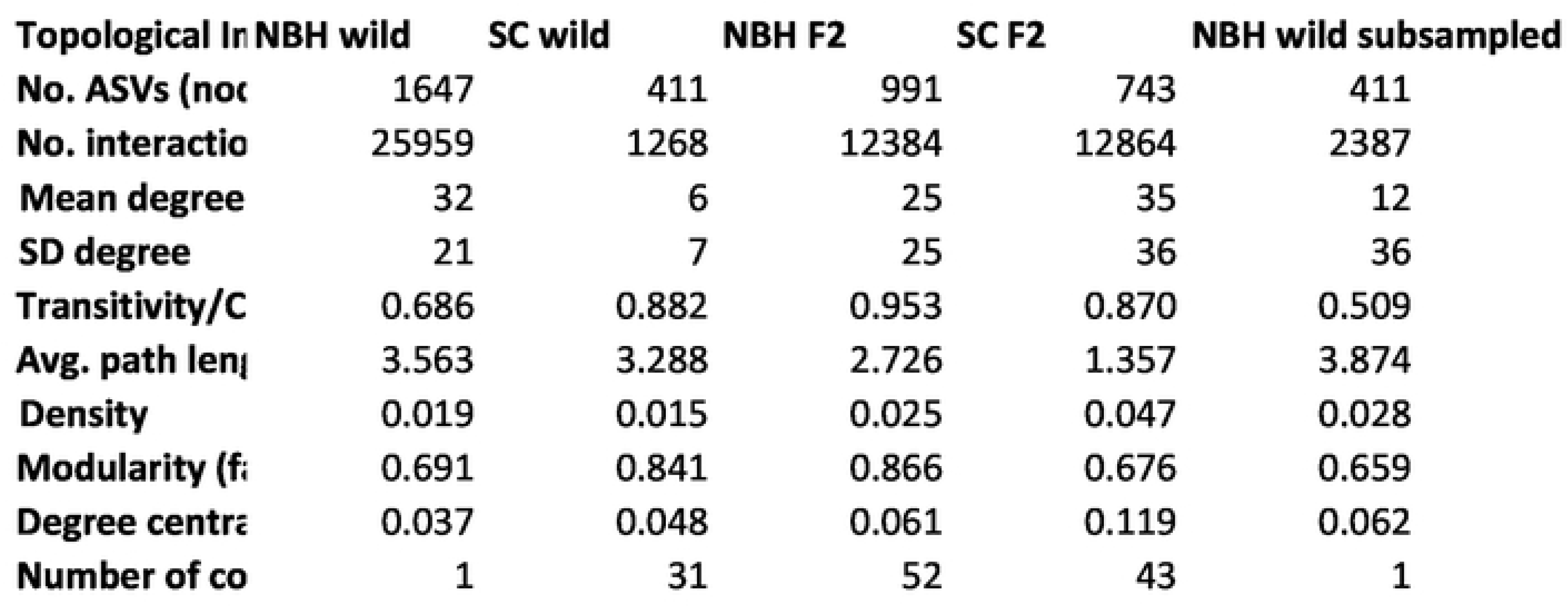
Mean alpha diversity metrics for all fish gut and water microbiomes. Numbers in parenthesis indicate the number of samples in each category.

**Table S2.**
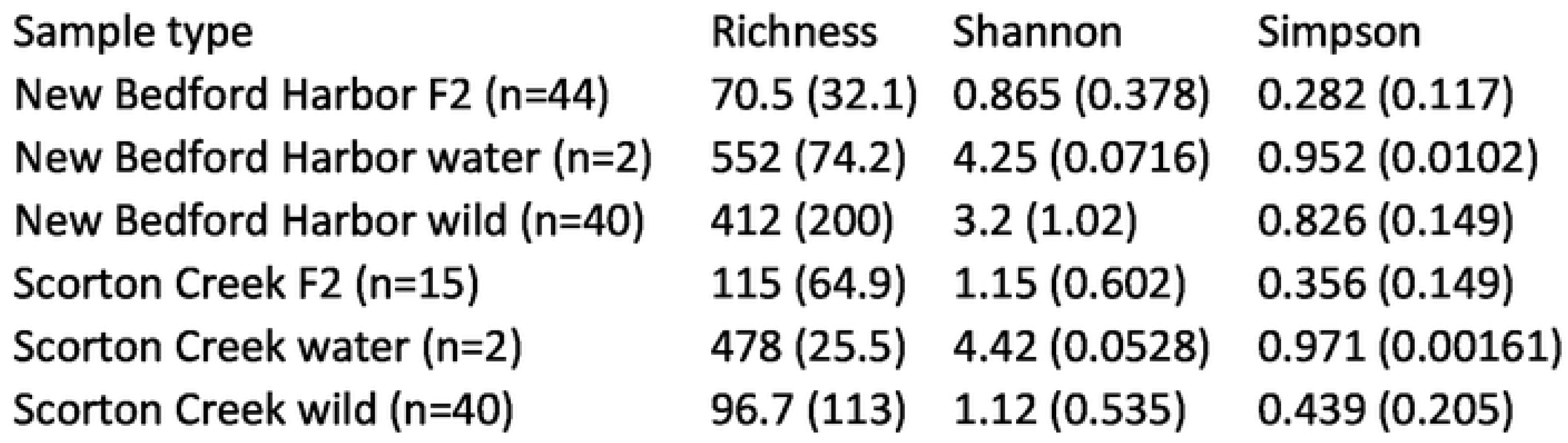
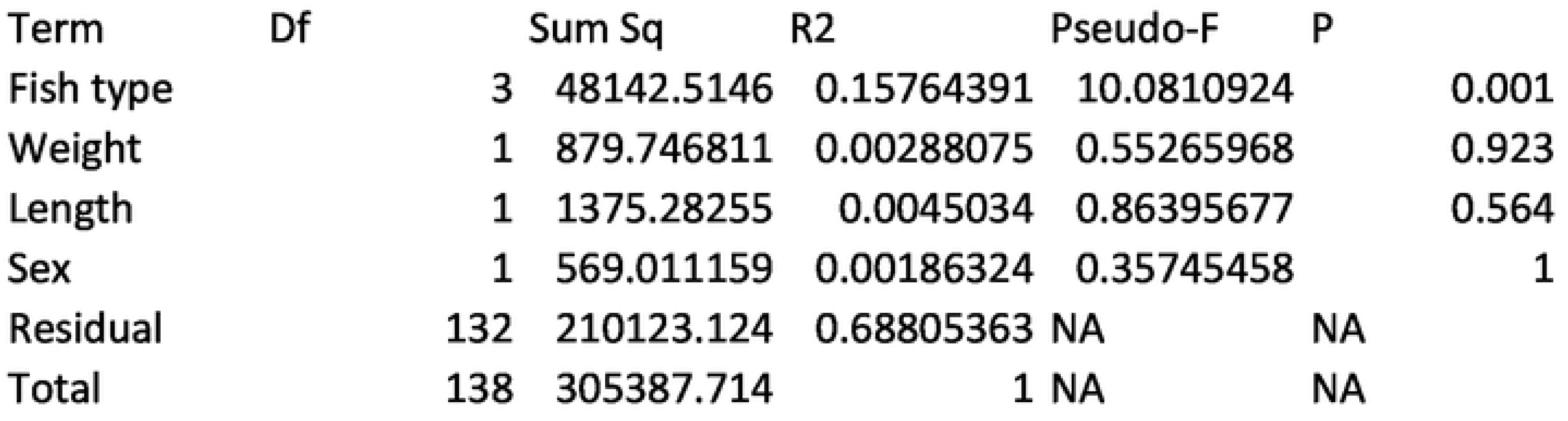
Marginal PERMANOVA on fish body condition.

**Table S3.**
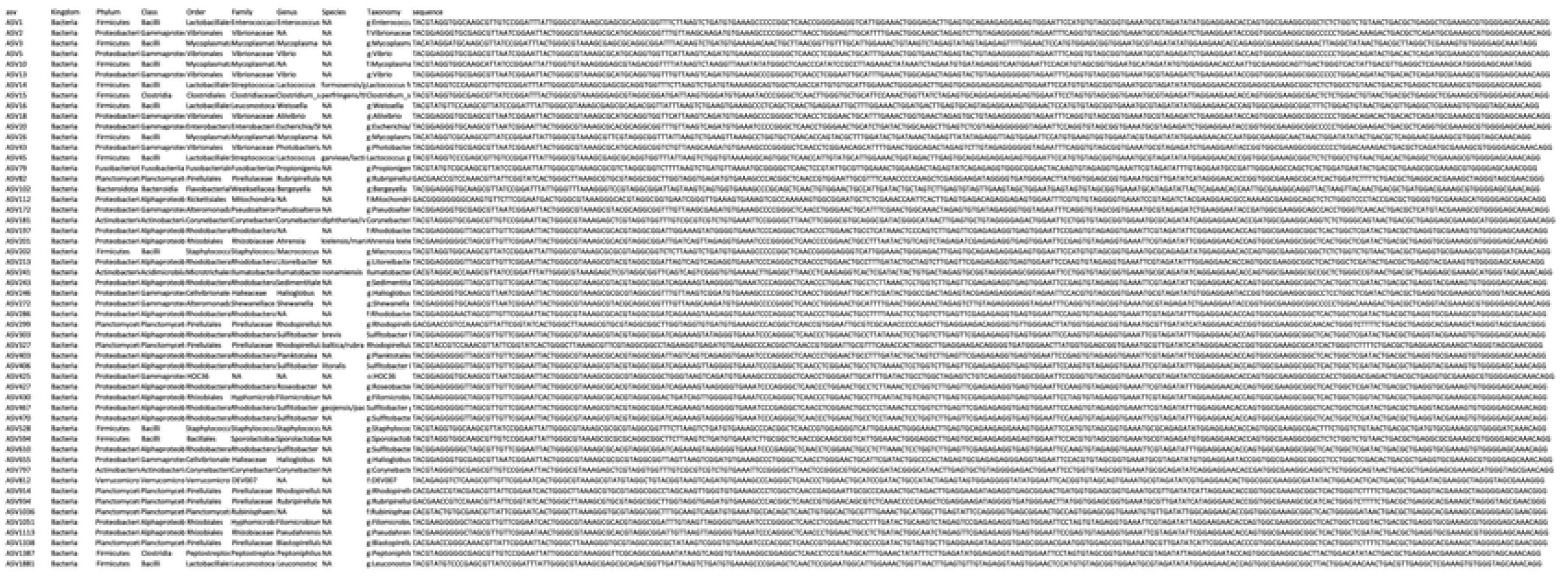
Taxonomic identity of shared ASVs between all fish types.

**Table S4.**
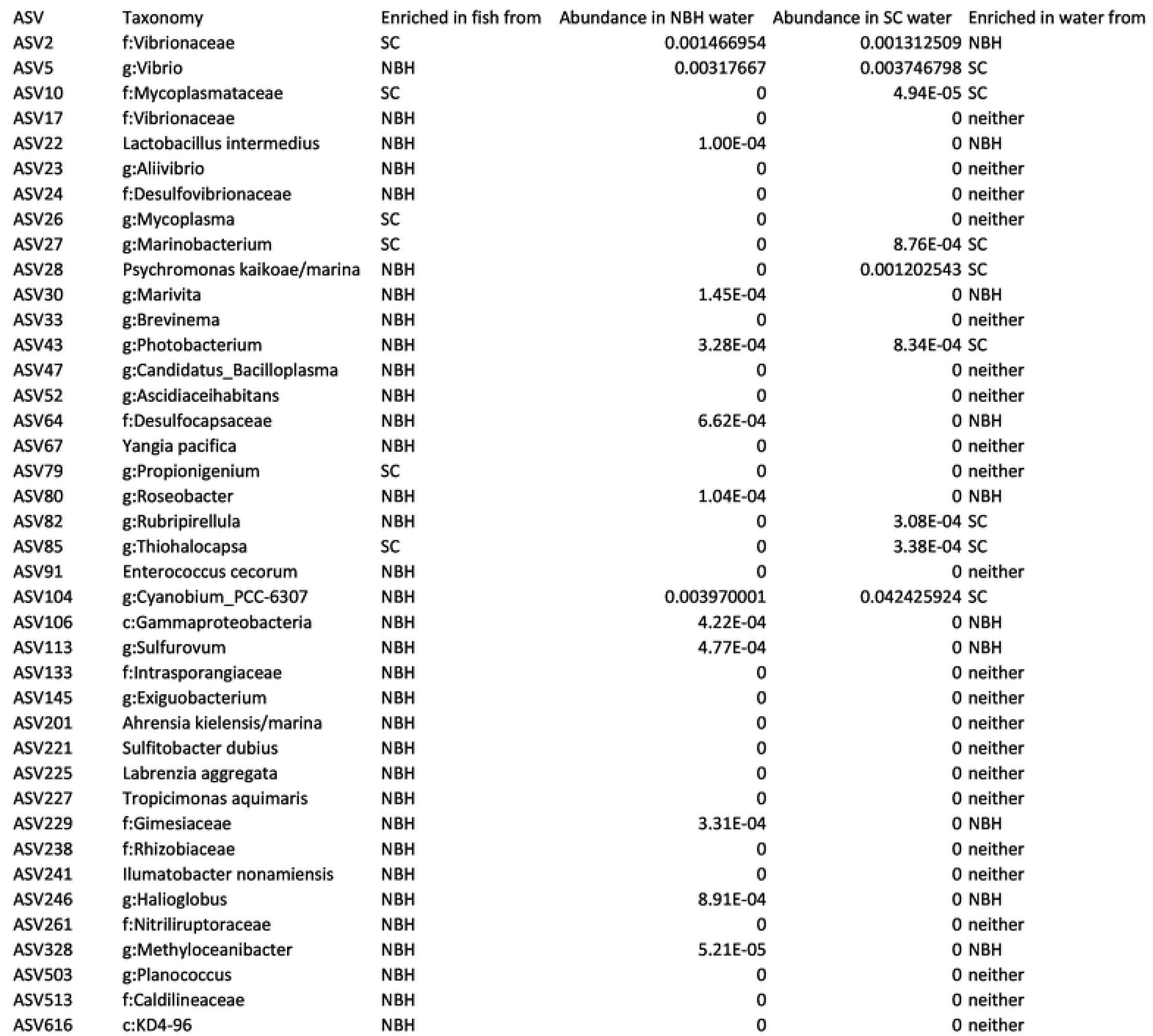
Abundance (in seawater samples) of ASVs calculated as significantly enriched between New Bedford Harbor wild and Scorton Creek wild fish microbiomes. The differential abundance of these ASVs in the fish microbiomes is not well explained by differing abundance in their aquatic environment.

**Table S5.**
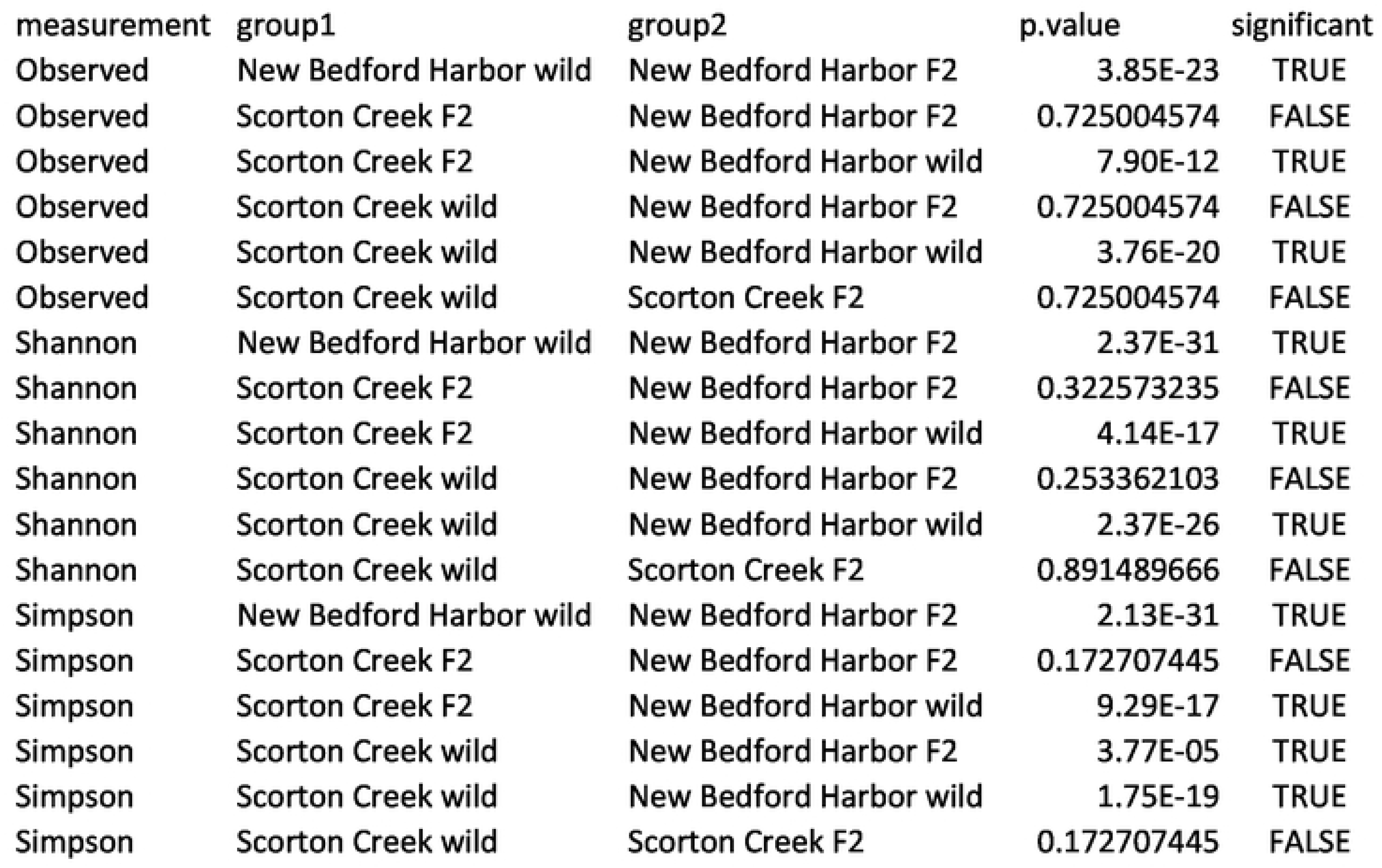
Pairwise t-tests of microbial community alpha diversity metrics between groups of fish, with holm correction for multiple comparisons.

**Figure S1.**
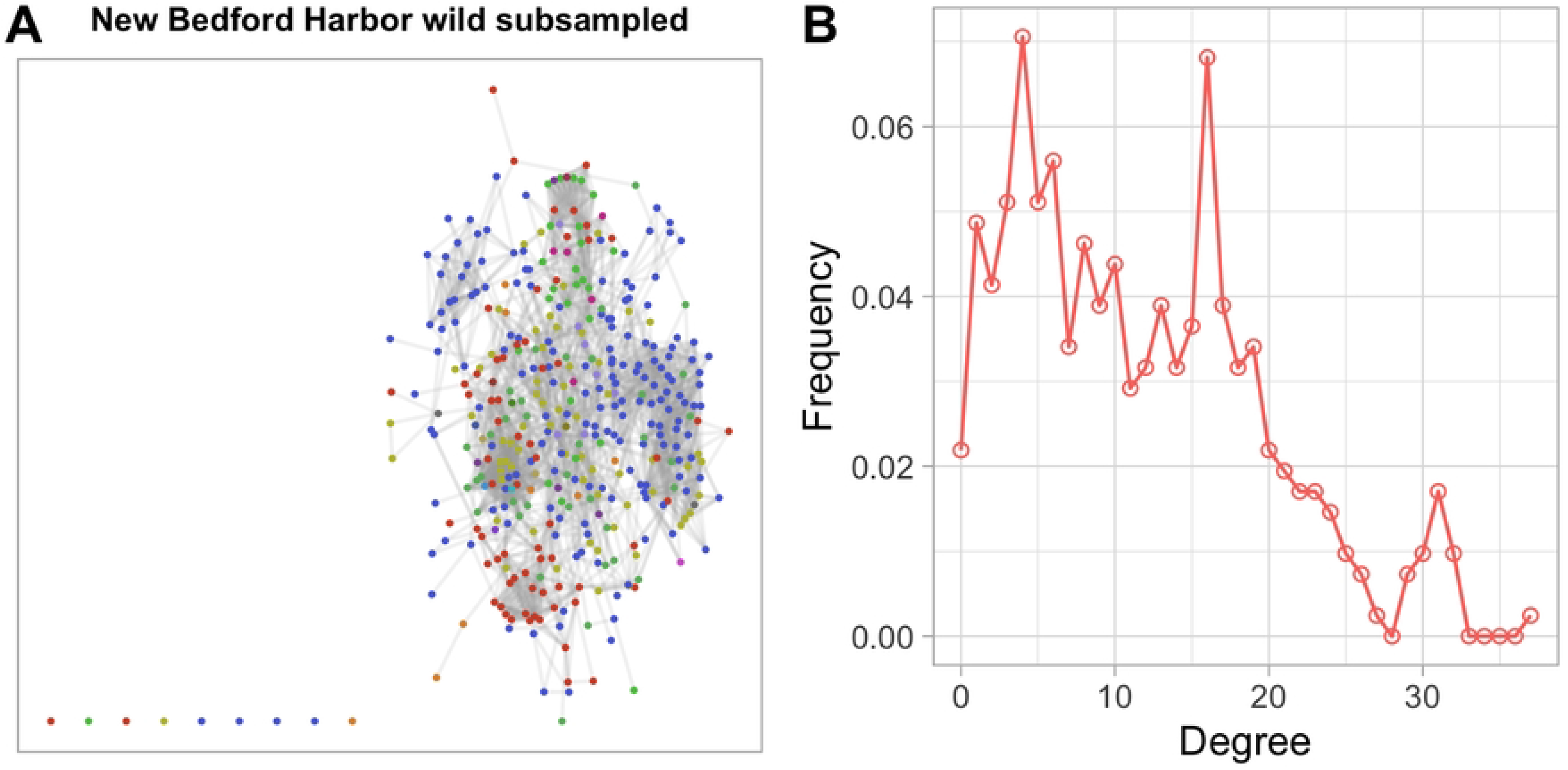
The subsampled microbial interaction network of New Bedford Harbor wild fish. (A) Network inferred after subsampling to the top 411 taxa to match the number of taxa in the Scorton Creek wild network. (B) Degree distribution of the subsampled network, showing similar pattern to the original network.

**Figure S2.**
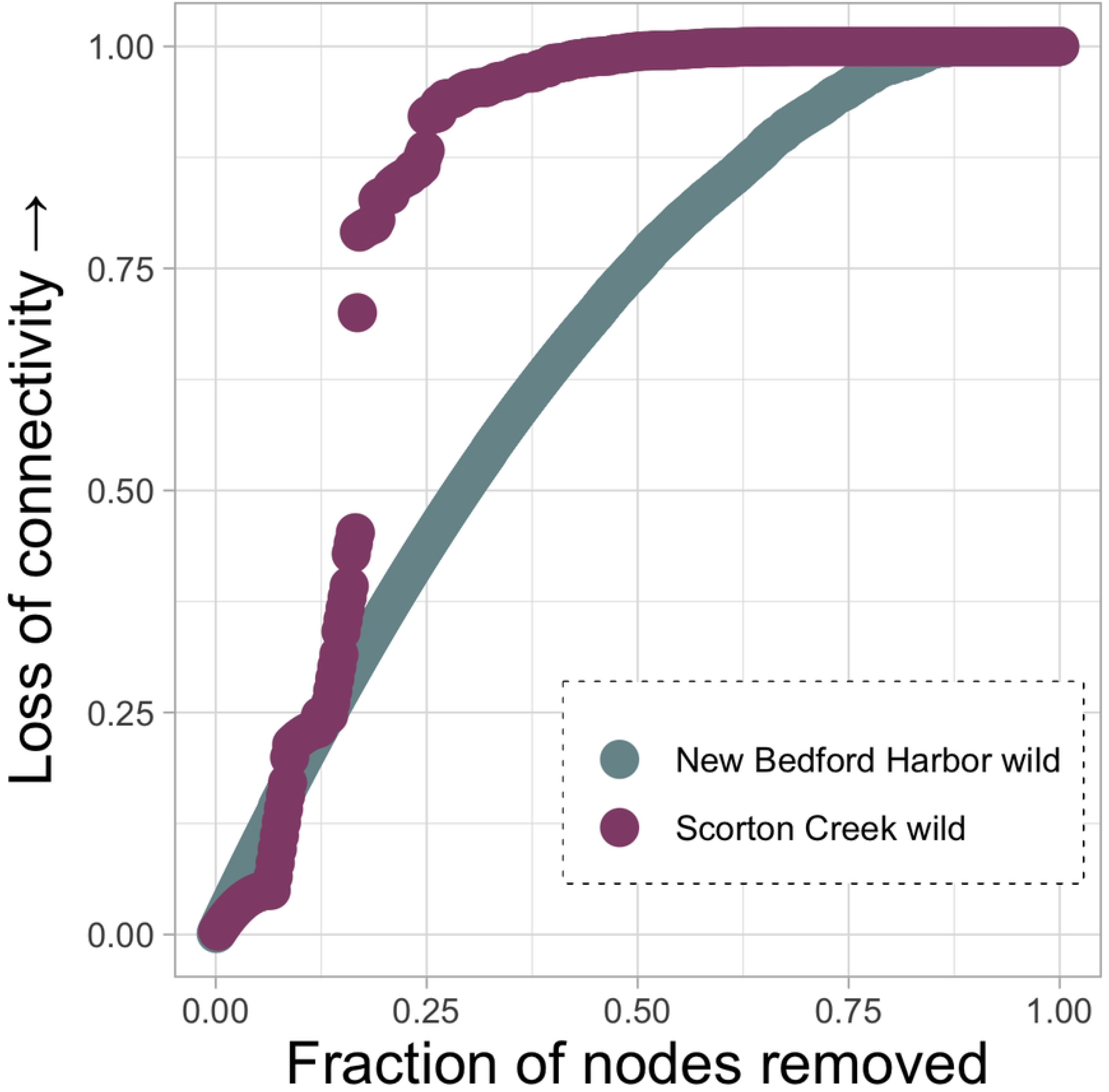
Sequential node removal more rapidly degrades the connectivity of SC wild network compared to NBH wild network. Nodes were removed in order of highest degree from both networks and the network connectivity loss as a percent of the original recalculated after each removal. Connectivity is measured as the sum of non-infinite shortest paths between nodes.

